# The rise and fall of the new variant of *Chlamydia trachomatis* in Sweden: mathematical modelling study

**DOI:** 10.1101/572107

**Authors:** Joost Smid, Christian L. Althaus, Nicola Low, Magnus Unemo, Björn Herrmann

## Abstract

**Objectives:** A new variant of *Chlamydia trachomatis* (nvCT) was discovered in Sweden in 2006. The nvCT has a plasmid deletion, which escaped detection by two nucleic acid amplification tests (Abbott/Roche, AR), which were used in 14 of 21 Swedish counties. The objectives of this study were to assess when and where nvCT emerged in Sweden, the proportion of nvCT in each county, and the role of a potential fitness difference between nvCT and co-circulating wild-type strains (wtCT).

**Methods:** We used a compartmental mathematical model describing the spatial and temporal spread of nvCT and wtCT. We parameterised the model using sexual behaviour data and Swedish spatial and demographic data. We used Bayesian inference to fit the model to surveillance data about reported diagnoses of chlamydia infection in each county and data from four counties that assessed the proportion of nvCT in multiple years.

**Results:** Model results indicated that nvCT emerged in central Sweden (Dalarna, Gävleborg, Västernorrland), reaching a proportion of 1% of prevalent CT infections in late 2002 or early 2003. The diagnostic selective advantage enabled rapid spread of nvCT in the presence of high treatment rates. After detection, the proportion of nvCT decreased from 30-70% in AR counties and 5-20% in counties that Becton Dickinson tests, to around 5% in 2015 in all counties. The decrease in nvCT was consistent with an estimated fitness cost of around 5% in transmissibility or 17% reduction in infectious duration.

**Conclusions:** We reconstructed the course of a natural experiment in which a mutant strain of *C. trachomatis* spread across Sweden. Our modelling study provides support, for the first time, of a reduced transmissibility or infectious duration of nvCT. This mathematical model improved our understanding of the first nvCT epidemic in Sweden and can be adapted to investigate the impact of future diagnostic escape mutants.

**Key messages:**

- The dynamics of a new variant of *Chlamydia trachomatis* (nvCT) that escaped testing and treatment in Sweden can be reconstructed using a mathematical transmission model.
- Our study for the first time provides support of a reduced transmissibility or infectious duration of the nvCT in Sweden.
- This mathematical model improved our understanding of the nvCT epidemic in Sweden and can be adapted to investigate the impact of future diagnostic escape mutants.

## INTRODUCTION

A genetic variant of *Chlamydia trachomatis* was discovered in Sweden in mid-September 2006.^1^ This variant, now referred to as the new variant of *C. trachomatis* (nvCT), has a 377 base pair deletion in the cryptic plasmid,^2^ which was part of the target sequence used by two nucleic acid amplification tests (NAATs). The deleted sequence resulted in thousands of false negative test results^3^ and nvCT spread undetected in 14 of 21 Swedish counties that used the Abbott m2000 (Abbott Laboratories, Abbott Park, IL, USA) and Cobas Amplicor/TaqMan 48 (Roche, Branchburg, NJ, USA) NAATs. The remaining seven Swedish counties used another NAAT, ProbeTEC (Becton Dickinson, BD, Franklin Lakes, NJ, USA) that used a different DNA target sequence on the plasmid. At the time of its detection in 2006-2007, the proportion of nvCT was between 20% and 65% in counties using the Abbott/Roche (AR) NAATs, compared with 7%-19% in counties using the BD NAAT.^3^

Testing of archived specimens found the earliest evidence of nvCT in Örebro (central Sweden) in one of 61 specimens in June 2003,^4^ but not in specimens from 1999-2000 in Örebro and from 2000-2001 in Malmö in southern Sweden.^5^ Statistical analyses of trends in CT positivity of specimens from Örebro from 1999 to 2006 indicated that nvCT increased to non-negligible levels between 2001-2002 and 2003-2004.^6^ There is little information about the presence of nvCT in other counties before 2006/2007. After the replacement of NAATs with versions that detect nvCT in AR counties, the proportion of nvCT had decreased to 5%-7% by 2015 in all counties, irrespective of the NAAT used.^7^

The studies published so far leave open questions about the dynamics of the emergence, spread and decline of nvCT. The timing and place of its emergence remain unknown. It is assumed that a single genetic event produced the new variant, which initially co-existed with the wild-type CT (wtCT) and then spread clonally, evading detection in areas that used AR tests.^5^ Extensive genomic comparisons and analyses of morphology, *in vitro* growth kinetics, phenotypic characteristics including antimicrobial susceptibility, and cell tropism did not detect any differences in fitness characteristics between any examined wtCT strain and nvCT.^5^ At the population level, the fitness of a CT strain can be expressed as the average number of secondary infections caused by an infected individual, i.e., the basic reproduction number *R*_0_.^8^ Hence, differences in the relative transmissibility or infectious duration of a strain would result in a either a fitness cost or benefit.^9^ So far, the available data could not determine the relative contributions of potentially altered transmission, patterns of sexual mixing between people infected with wtCT and nvCT, and changes in testing to the observed nationwide increase and subsequent decrease in the proportion of nvCT.^10^

The objectives of this study were to assess when and where nvCT emerged in Sweden, the proportion of nvCT in each county, and the role of a potential fitness difference between nvCT and the co-circulating wtCT strains. To this end, we developed a mathematical model of CT transmission that was fitted to surveillance data about reported diagnoses of chlamydia infection in each county, and data from four counties that assessed the proportion of nvCT in multiple years, using Bayesian inference.

## METHODS

### Data

We used data on the type of NAAT used in Swedish counties. From the mid-1990s, chlamydia cell culture was replaced by NAAT. In 13 of 21 Swedish counties, AR NAATs were used, in seven counties, the BD NAAT system was used and one county, Västra Götaland, used the BD NAAT in three of four laboratories and an AR NAAT in a single smaller laboratory. We refer to counties according to the NAAT system used, with the suffix ‒AR or ‒BD. In the model, Västra Götaland is treated as a BD county.

We used published data about the proportion of nvCT and data about diagnosis rates in different years and counties in Sweden. Data about the proportion of nvCT were published for seven AR counties and five BD counties just after its discovery (late 2006 and early 2007).^3^ Additional data for 2008, 2009, 2011 and 2015 were available for four of these 12 counties (Dalarna-AR, Örebro-AR, Uppsala-BD and Norrbotten-BD, Table S1).^7^ We had data about the yearly number of CT diagnoses from all counties since 1997.^11^ In the model, we only used CT diagnosis data from Dalarna, Örebro, Uppsala and Norrbotten for selected years before and after the discovery of nvCT (2004, 2006, 2007, 2008 and 2009, Table S2). We chose this subset of data to give the diagnoses rates and the data about the proportion of nvCT a similar weight for Bayesian inference. We also anticipated that those differences in diagnosis rates between AR counties and BD counties, attributable to misdiagnoses by AR NAAT before the discovery of nvCT, would be most pronounced in years around its discovery. For simplicity, we assumed that only people aged 15-29 years old were tested. We calculated the diagnosis rate for this population by dividing the total number of diagnoses by the county-specific yearly population sizes for this age group. Population sizes were obtained from Statistics Sweden.^12^

We used spatial data specifying distances between the centroids of the counties, which we downloaded as a geographic information system shapefile. Lastly, we used sexual behaviour data about the yearly number of new heterosexual partners for sexually experienced people from 15 to 29 years. These data were from the second British National Survey of Sexual Attitudes and Lifestyles (Natsal-2) because there were no detailed sexual behaviour data from Sweden. Natsal-2 is a nationally representative cross-sectional survey that interviewed 12,110 respondents aged 16-44 years (response rate 65.4%) from 1999 to 2001.^13^ The data collection period is close to the likely timing of nvCT emergence. We did not include data from Natsal-3 (2010-2012) as changes in sexual behaviour between the two surveys were small.^14^

### Transmission model

We developed a mathematical model to describe the spread of nvCT in Sweden (table 1, Supplementary Material S1). We implemented the spatial structure of Sweden consisting of 21 counties using a gravity model in a meta-population framework.^15–17^ We considered the population of 15-29 year old Swedish sexually experienced heterosexual adults, subdivided into a low and a high sexual activity class (figure S1). Individuals can be infected with either wtCT or nvCT. We assumed that wtCT is representative of all CT strains that are co-circulating with nvCT. Susceptible people seek sexual partners at rate *c*. The per partnership transmission probability of wtCT and nvCT is *β* and *fβ*, respectively, where *f* can account for a potential difference in transmissibility of nvCT. Infected people clear infection naturally at rate *γ*, or receive treatment.

**Table 1.**
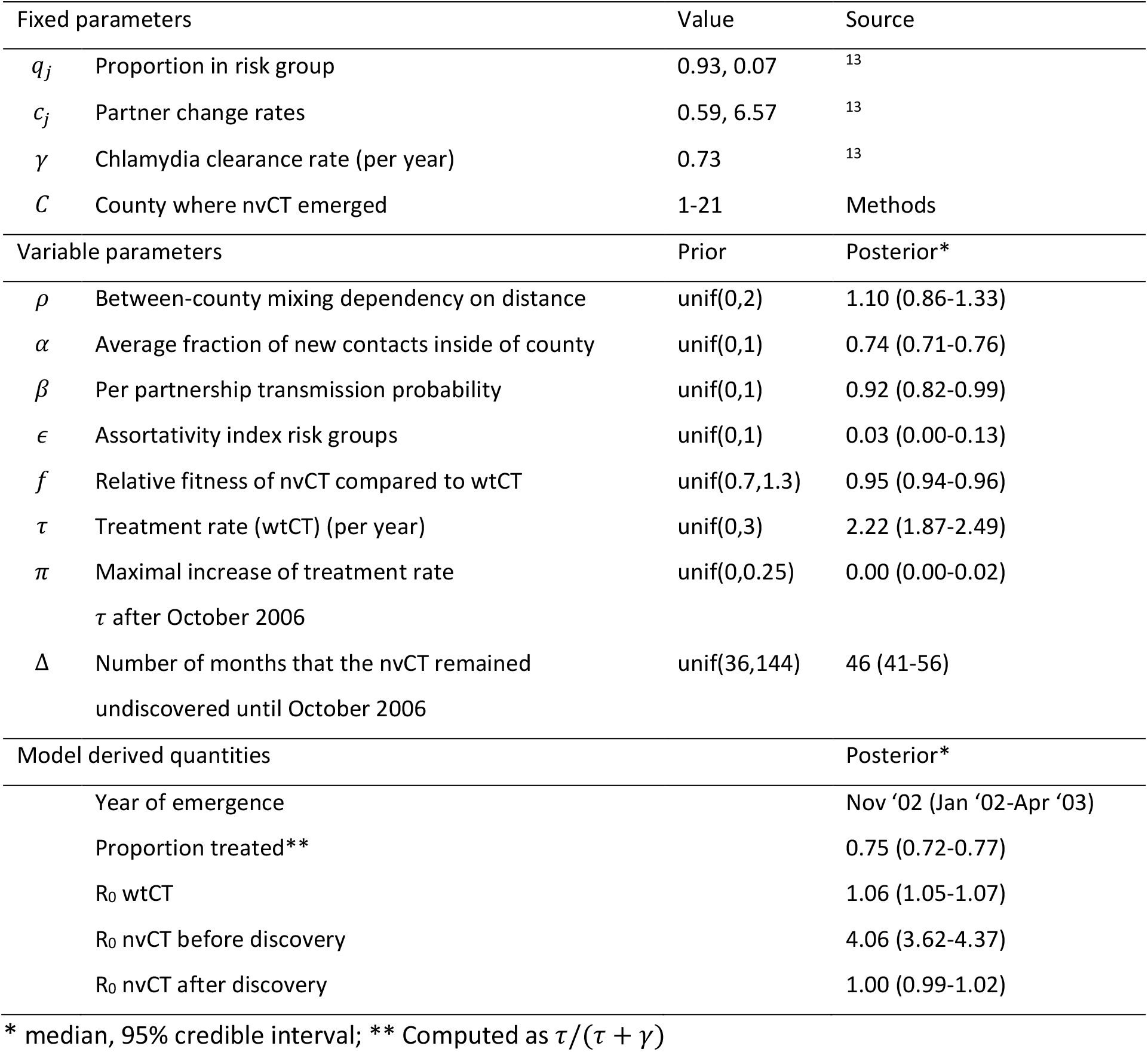
Description of fixed and variable model parameters, and model derived quantities. Only results for the model in which we assumed that nvCT emerged in Dalarna are shown.

We assumed that wtCT infected people in all counties are treated at a fixed rate *τ* before October 2006. nvCT infected people are treated at the same rate in BD counties, whereas they were not treated in AR counties before 2006. We assumed that after October 2006, when nvCT was discovered and the AR tests were modified, all people with diagnosed CT were treated. The increased number of tests after 2006 (Supplementary Material, figure S2) and reported diagnosis rates could be a result of a higher treatment rate directly after the discovery of nvCT.^10^ We modelled this by assuming that the treatment rate is a certain percentage (*π*) higher than *τ* directly after the discovery of nvCT, and that this percentage decreases linearly to zero within 3 years.

### Bayesian inference

We obtained posterior distributions of the model parameters by fitting the model to two data sets using Bayesian inference (Supplementary Material). To determine the most likely origin of nvCT emergence, we analysed 21 versions of the model; in each version, the county in which nvCT emerged differed. We then used the deviance information criterion (DIC)^18 19^ to compare the model fits.

### Sensitivity analyses

We investigated the alternative assumption that a fitness difference between wtCT and nvCT strains is associated with a relative difference in infectious duration instead of a relative difference in transmissibility.

## RESULTS

The model version in which nvCT emerged in Dalarna-AR (central-Sweden) fitted best to the empirical data, as indicated by the lowest DIC value (figure 1). The next best fitting models assumed the emergence of nvCT in Gävleborg-AR and Västernorrland-AR (also in central Sweden). The remainder of the results are from the model assuming emergence in Dalarna.

**Figure 1.**
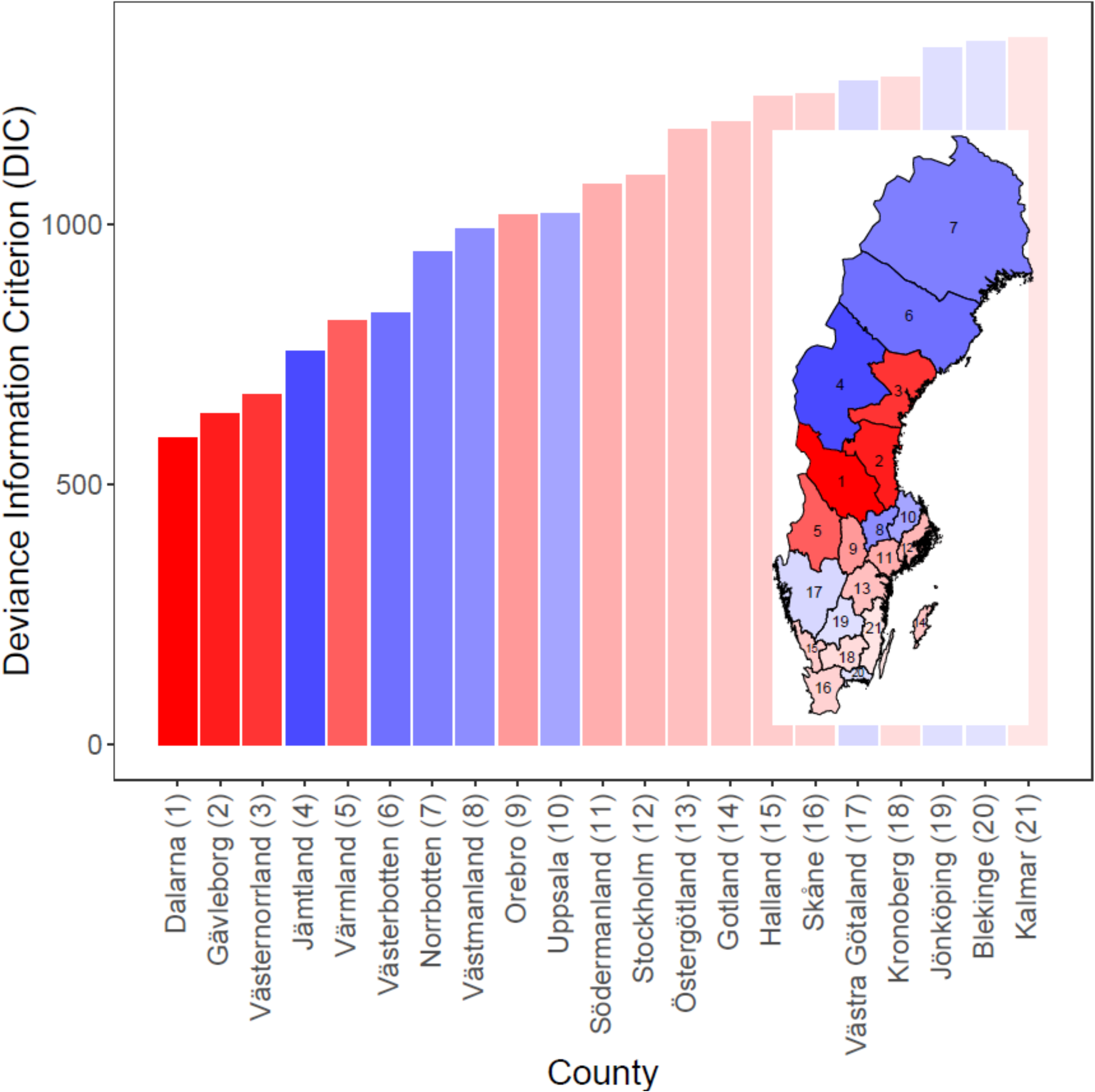
Deviance information criterion (DIC) values for 21 different models, each of which assumes that nvCT emerged in that county. Red bars are Abbott-Roche (AR) counties, blue bars are Becton Dickinson (BD) counties. Intensity of shading inversely proportional to DIC.

Overall, the model fitted well to the available empirical data about the proportion of nvCT (figure 2). For the 12 counties with data just after the discovery of nvCT in late 2006 and early 2007, the proportion of nvCT was higher in the AR counties than in the BD counties (figure 2A). The proportion of nvCT in Västra Götaland was less well captured by the model, which might result from the mix of BD and AR tests used in different laboratories. For the four counties with more frequent data, the proportion of nvCT was higher in Dalarna-AR and Örebro-AR than in Uppsala-BD and Norrbotten-BD in 2006-2007, after nvCT was discovered (figure 2B). By 2015, the proportion of nvCT converged to around 5% in all four counties. The model also captures trends in diagnoses rates, although there are some discrepancies for specific years and counties (figure 2C). In particular, the predicted diagnosis rates are lower than the actual diagnoses rates for Dalarna-AR after the discovery of nvCT, and for Örebro-AR that was the case for 2008 and 2009.

**Figure 2.**
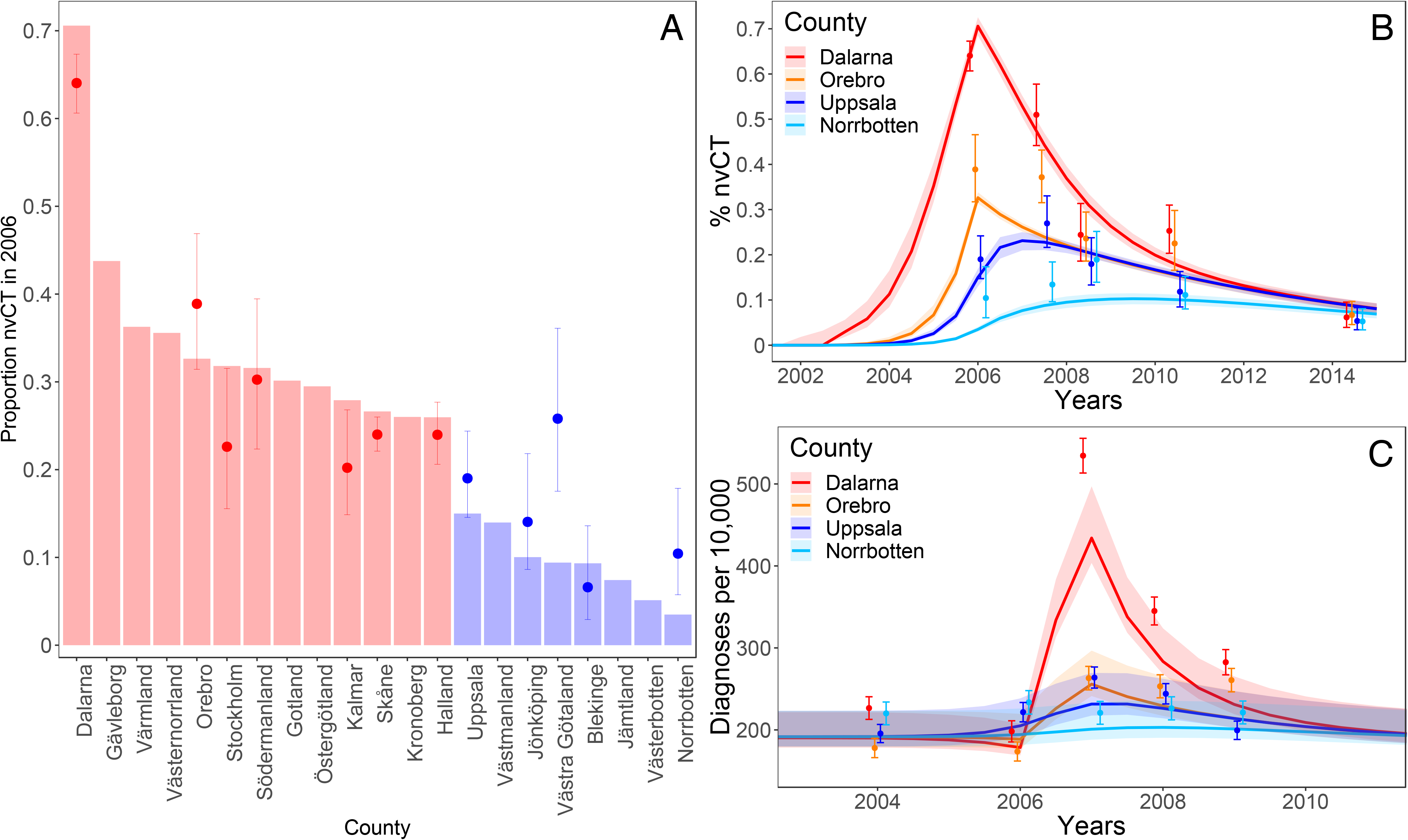
Data and model fit for proportions of nvCT in all counties in 2006-2007, after the discovery of nvCT (A), proportions of nvCT in Dalarna, Örebro, Uppsala and Norrbotten between 2000-2015 (B), and number of CT diagnoses per 100,000 population in Dalarna, Örebro, Uppsala and Norrbotten between 2000-2015 (C). Dots and error bars: median values and 95% confidence intervals for the data. Bars (A) and trajectories (B, C): model outputs. Red bars are Abbott-Roche (AR) counties, blue bars are Becton Dickinson (BD) counties.

In our model, the posterior distribution of the parameter ∆ indicates that nvCT remained undetected for at least 46 months (Bayesian credible interval (BCI): 41-56 months) after its introduction until its discovery in October 2006 (table 1). Using this estimate, nvCT represented 1% of prevalent infections in Dalarna in November 2002 (BCI: January 2002-April 2003) (figure 3). By the time that nvCT was discovered in late 2006, the model suggests that nvCT had spread to all counties, with the highest proportions of nvCT in AR counties (figures 2A and 3). The rapid growth was primarily driven by the high treatment rate in the model, i.e., the selective advantage of nvCT of not being treated in AR counties. The model-estimated treatment rate (*τ*) was 2.22 (BCI: 1.87-2.49) treatments per infected person per year. Infected people can either clear infection naturally at rate *γ* or be treated at rate *τ*. Therefore, using the equation *τ*/(*τ* + *γ*), we estimated that 75% (BCI: 72-77%) of the wtCT infected population was treated.

**Figure 3.**
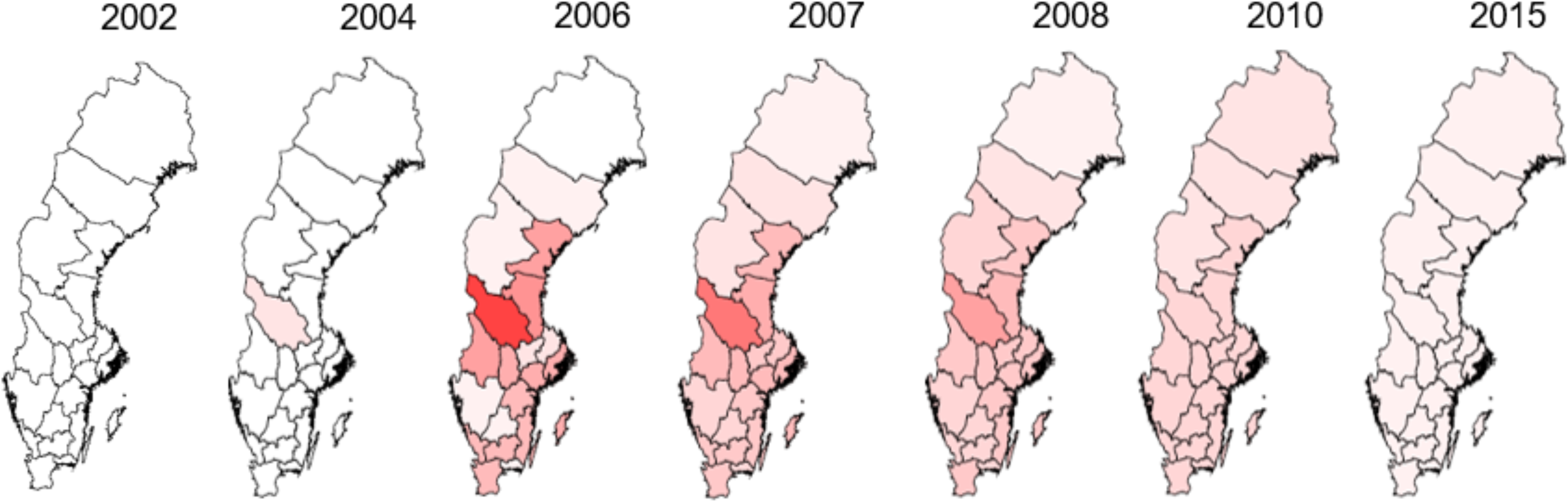
Rise and fall of nvCT between 2002 and 2015 in different Swedish counties. The shading indicates the proportions of nvCT cases among all CT cases.

The observed CT diagnosis rates in AR counties were much higher after the discovery of nvCT than before (figure 2C). In the model, this observation reflects increased chlamydia prevalence after emergence of nvCT because people in AR counties with undiagnosed nvCT did not receive treatment until 2006. The parameter *π* (increase in testing rate *τ* from 2006-2009, after discovery of nvCT) in the model represents increased testing. In the model, *π* was estimated at close to zero, so increased transmission alone could account for the increase in diagnosis rates. In reality, the increase in testing that occurred in Sweden in 2007, might also have contributed to this increase.

From 2007, the proportion of nvCT and the diagnosis rates decreased in all counties. We estimated that nvCT was 5% (BCI: 4-6%) less transmissible than wtCT strains. In the sensitivity analysis where we assumed that the fitness cost could also be associated with infectious duration, we estimated a 17% (BCI: 16-19%) reduction in infectious duration for nvCT compared to wtCT strains (Supplementary Material, figure S4). In both cases, the reduced fitness of nvCT and its dilution across counties, together with the loss of the diagnostic selective advantage, led to the decline in its proportion after its discovery. This process can also be quantified by *R*_0_, which we computed for wtCT and nvCT in presence and absence of effective testing and treatment (Supplementary Material, S3). *R*_0_ for wtCT is slightly above the threshold of one (1.06, BCI: 1.04-1.06). *R*_0_ for nvCT before its discovery is considerably higher (4.06, BCI: 3.62-4.37), owing to the diagnostic selective advantage of nvCT compared with wtCT. In contrast, *R*_0_ for nvCT after its discovery is lower (1.00, BCI: 0.99-1.02), owing to its reduced transmissibility and the lost diagnostic selective advantage.

## DISCUSSION

Using a mathematical model we were able to reconstruct the course of a natural experiment in which a mutant strain of *C. trachomatis* escaped detection and spread across Sweden. According to our model, the nvCT emerged in central Sweden (Dalarna, Gävleborg, Västernorrland), reaching a proportion of 1% of prevalent CT infections in late 2002 or early 2003. The diagnostic selective advantage of nvCT enabled its rapid spread in the presence of high CT treatment and transmission rates. After detection, the proportion of nvCT decreased from 30-70% in AR counties and 5-20% in BD counties to around 5% in 2015 in all counties because of between-county mixing and the subsequent dilution of nvCT. Our modelling results suggest that this decline is consistent with the lost diagnostic selective advantage and reduced fitness of the nvCT, either because of reduced transmissibility by around 5%, or reduced infectious duration by around 17%. Our modelling study for the first time provides support of a reduced transmissibility or infectious duration of nvCT.

The strengths of the model are that it is relatively simple, but incorporates necessary heterogeneity in mixing between groups with differing levels of sexual activity and between different geographical areas. There were sufficient empirical data about the proportion of nvCT after its discovery and more detailed data in both AR and BD counties to fit the model. Our study also has a number of limitations that need to be considered when interpreting the findings. First, we had to make some simplifying assumptions about the structure and parameters of the transmission model. Owing to the limits of the MCMC inference framework, we decided to use fixed values for the sexual partner change rates and the chlamydia clearance rate. There is considerable uncertainty with regards to the exact values of these parameters, which might explain the relatively high posterior values of the per-partnership transmission probability, and the finding that sexual mixing between activity classes is close to random. However, the differential spread of wtCT and nvCT is mainly driven by the detection/treatment rate and the fitness cost and we previously showed that the transmission probability and mixing coefficient only marginally influence the relative spread of different strains of sexually transmitted infections (STIs).^9^ The relatively simple model structure makes the modelled prevalence sensitive to small changes in the transmission dynamics. While our model can describe changes in the proportion of nvCT well, it does not necessarily generate reliable estimates of incidence and prevalence. Second, we did not model the possibility that factors other than the spread of nvCT, such as increased incidence or changes in screening policy, might have contributed to an increase in overall CT prevalence in Sweden.^20^ Third, we used a deterministic model. While very unlikely, nvCT could have emerged independently in multiple unrelated people, possibly in different counties and at different times. Modelling these possibilities in more detail would require a stochastic modelling framework. The choice of a deterministic model did not affect parameter inference, because we only used data for years when there was a substantial proportion of nvCT. Modelling stochasticity in these circumstances is less important.^15^ Furthermore, extensive genotyping of nvCT cases from different time points and countries support a clonal spread of nvCT.^3 7 21 22^ Finally, we did not include the possibility of acquired immunity in our model. More untreated infections in AB counties could have resulted in higher levels of immunity in the population. It is unclear how this would have affected the differential spread of nvCT and wtCT.

The emergence of nvCT in Sweden was, to our knowledge, the first reported example of a strain mutating to escape detection by a diagnostic test, but was not unique. In May 2019, researchers in Finland reported false-negative chlamydia test results using the Aptima Combo 2 test, resulting from a single point mutation in the 23s rRNA gene.^23 24^ These events in the field of diagnostics can be compared to the phenomenon of immune escape, where viral pathogens such as human immunodeficiency virus (HIV) evade recognition by cytotoxic T lymphocytes through mutation in their epitopes.^25^ The emergence of nvCT is also related to the mechanisms that facilitate the spread of antimicrobial resistance for other STIs. High treatment rates are the major driver for the spread of *Neisseria gonorrhoeae* strains that are resistant to antimicrobials such as ciprofloxacin.^9,26^ Antimicrobial resistance in CT has not been detected yet, but its possible emergence cannot be excluded with continued antimicrobial selection pressure.^27^ Our estimate that around three quarters of infected people receive treatment (Supplementary Material, S5) is consistent with the history of early introduction of widespread opportunistic testing and an advanced system of partner notification that identifies recent sex partners with a high probability of CT.^28 29^ The insights into the means by which competition between CT strains, through selection pressure induced by testing and treatment, can inform the future study of the potential for the spread of antimicrobial resistance in CT. Increased levels of STI testing with molecular diagnostic methods mean that mutations that affect the target sequences for nucleic acid amplification provide new evolutionary advantages for *C. trachomatis* and other pathogens.

This modelling study provides insights into the spread of nvCT in Sweden that are consistent with the evidence obtained from epidemiological and genomic studies. Our study shows the value of high quality surveillance data and of the prompt investigation and follow up of anomalies in these data. The estimated year of emergence agrees with retrospective analyses that first detected nvCT in 2003,^4^ with presumed emergence between 2001-2002 and 2003-2004^6^ and absence of detection of nvCT before 2002.^5^ In addition, our study indicates that nvCT was associated with a fitness cost at the population level, which resulted in reduced transmission. *In vitro* studies showed that nvCT had unaltered fitness compared with all examined wtCT strains.^5^ However, small-scale gene variations and/or alterations in timing and level of gene transcription and translation affecting the fitness *in vivo* or *in vitro*, could not be excluded.^5^ Also, fitness as measured by *in vitro* growth assays is not necessarily related to fitness at the population level, as expressed by *R*_0_. In conclusion, our model results quantified the speed of spread of *C. trachomatis* in the absence of testing and treatment.

This mathematical model improved our understanding of the first nvCT epidemic in Sweden and can be adapted to investigate the impact of future diagnostic escape mutants.

## Acknowledgements

None.

## Contributors

BH and NL designed and coordinated the study. JS with support from CLA and NL performed the modelling. All authors were involved in the analysis and interpretation of the results, and writing and approving of the final version of the manuscript.

