## Supplementary material for "The rise and fall of the new variant of *Chlamydia trachomatis* in Sweden: mathematical modelling study"

### S1: Model description

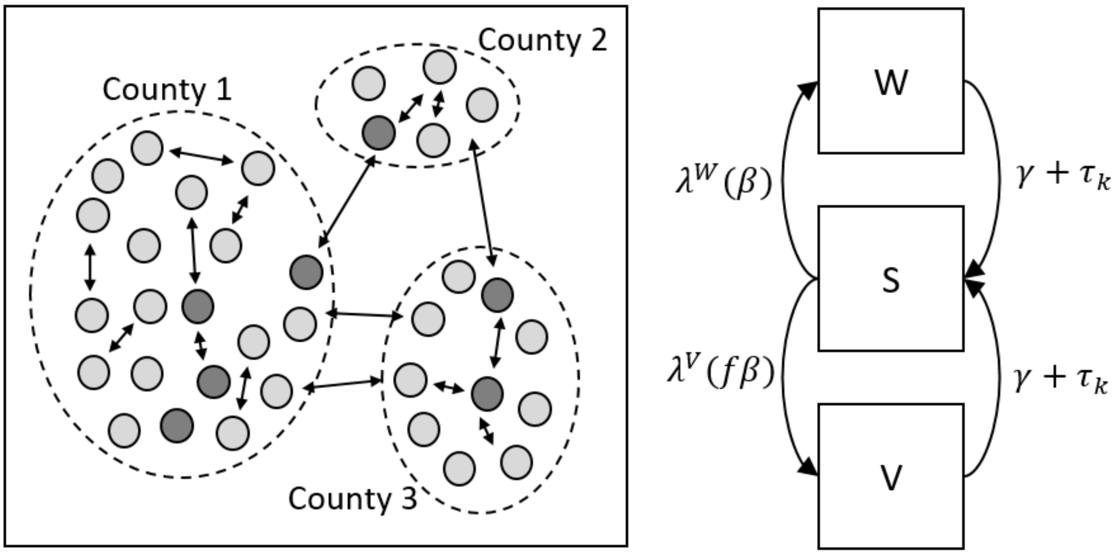

*Figure S1: A) Representation of a meta-population model for sexually active heterosexual people of lower sexual activity class (light grey) and of a higher sexual activity class (dark grey) in three different counties. The proportion of high-activity individuals is the same in all counties. Partnerships are represented as two-head arrows. (B) Structure of infection transmission model. Individuals have three infection states: uninfected and susceptible (S), infected with wtCT (W) and infected with nvCT (V). Arrows represent transitions between infection states. Transition rates: $\lambda$^$W(\beta)$ force of infection for becoming wtCT infected, depending on transmission probability $\beta$; $\lambda$^$V(f\beta)$ force of infection for becoming nvCT infected, depending on transmission probability $\beta$ and fitness difference for nvCT ($f$); $\gamma$ natural clearance rate; $\tau_k$ treatment rate, depending on type of county $k$ (AR/BD). All rates except $\gamma$ are also time-dependent.*

#### Overview

We developed a mathematical model to describe heterosexual *C. trachomatis* transmission and the spread of the nvCT in Sweden (Figure S1). We implemented the spatial structure of Sweden consisting of 21 counties as a meta-population model. We modeled the population of 15-29 year old Swedish sexually experienced adults, subdivided into a low and a high sexual activity class. In the model, people can be susceptible people ($S$), infected with the wtCT ($W$) and infected with the nvCT ($V$). We used the following system of ordinary differential equations (ODE):

$$\frac{dS_{j,k}}{dt} = -(\lambda_{jk}^W + \lambda_{jk}^V)\, S_{jk} + \gamma(W_{jk} + V_{jk}) + \tau_k(t)(W_{jk} + V_{jk})$$

$$\frac{dW_{jk}}{dt} = \lambda_{jk}^{W} S_{jk} - \gamma W_{jk} - \tau_k(t) W_{jk}$$

$$\frac{dV_{jk}}{dt} = \lambda_{jk}^{V} S_{jk} - \gamma V_{jk} - \tau_k(t) V_{jk}$$

Here subscripts $j$ denotes sexual activity class and $k$ the county. Susceptible people can be infected with the wtCT or the nvCT at rate $\lambda_{jk}^{W}$ and $\lambda_{jk}^{V}$, respectively (the forces of infection). Infected people clear infection naturally at rate $\gamma$, or may be treated for infection at rate $\tau_k(t)$, dependent on time, county and on whether they were infected with the wtCT or with the nvCT. We assume in the model that before October 2006, in all counties wtCT infected people were treated at a fixed rate $\tau$. In counties using AR tests, nvCT infected people were not treated, whereas nvCT infected people were treated at the same rate $\tau$ in counties using BD tests. We assume that after October 2006, all infected people were treated. The increased number of tests done directly after 2006 (Fig. S2) suggests a higher treatment rate after the nvCT was discovered. We model this by assuming that $\tau_k(t)$ was a certain percentage point ($\pi$) higher than $\tau$ in October 2006, and that this percentage linearly decreased to zero within 3 years after this month. Therefore, $\tau_k(t)$ is parameterized as follows:

$$\tau_k(t) = \begin{cases} \tau & if\ k = BD\ and\ t < 2007 \\ 0 & if\ k = AR\ and\ t < 2007 \\ \tau + \pi\tau(1 - 1/3(t - 2007)) & if\ k \in \{AR, BD\}\ and\ 2007 \leq t < 2010 \\ \tau & if\ k \in \{AR, BD\}\ and\ t \geq 2010 \end{cases}$$

### Sexual mixing and force of infection

In our model, the time-dependent forces of infection exerted by the wtCT ($\lambda_{jk}^{W}$) and nvCT ($\lambda_{jk}^{V}$) depend on assumptions about transmission rates and sexual contact preferences between individuals from different sexual activity classes and counties. We model them as:

$$\lambda_{jk}^{W} = c_j \beta \sum_i \sum_l M_{kl} (\epsilon \delta_{ij} + (1 - \epsilon) \frac{c_i N_{il}}{\sum_m c_m N_{ml}}) \frac{W_{il}}{N_{il}}$$

$$\lambda_{jk}^{V} = c_j f \beta \sum_i \sum_l M_{kl} (\epsilon \delta_{ij} + (1-\epsilon) \frac{c_i N_{il}}{\sum_m c_m N_{ml}}) \frac{V_{il}}{N_{il}}$$

Here $c_j$ is the sexual partner change rate and $\beta$ the per partnership transmission probability. Parameter $f$ denotes the relative fitness of the nvCT compared to the wtCT, where we assume that a potential fitness difference between the wtCT and the nvCT increases or decreases the per partnership transmission probability, rather than the duration of infection.

We model a certain degree of assortative mixing with respect to activity classes $i$ and $j$ by parameter $\epsilon$, which takes values between 0 and 1. Here, $\delta_{ij}$ is the Kronecker delta, which equals 1 if $i = j$ and 0 otherwise. Therefore, $\epsilon = 0$ corresponds to proportional (random) mixing where sexual partners are chosen in proportion to the size of their sexual activity class; $\epsilon = 1$ corresponds to fully assortative mixing where people only have sexual contacts with people from the same sexual activity class.

Sexual mixing between individuals from counties $k$ and $l$ was modeled through matrix $M_{kl}$, containing the conditional probabilities that somebody from county $k$ has a sexual contact with somebody from county $l$. We assumed a gravity model, i.e., mixing between counties is positively associated with the population size and inversely related to the distance between counties. Owing to the higher partner change rate, individuals from the high sexual activity class will be more likely to engage in between-county partnerships than individuals from the low sexual activity class. We defined the gravity matrix $\Phi$ with entries

$$\Phi_{kl} = \frac{N_k N_l}{d_{kl}^{\rho}}$$

where $N_k$ is the population size of county $k$, and $d_{kl}$ is the distance between the geographical centers of counties $k$ and $l$, as a proxy for the average distance of all contacts in these counties. Parameter $\rho$ controls the dependence of sexual mixing on distance between people living in different counties. Using $\Phi$ only, mixing within counties as compared to mixing between counties is still undefined. Therefore, we used algebraic manipulations proposed by Riesen et al.[1] to reconstruct the mixing matrix $M$ from $\Phi$. We rescaled $\Phi$ by a scaling factor $s$ and weighted all columns with the inverse of the population size of a county:

$$M_{kl} = s \frac{\Phi_{kl}}{N_k}$$

To transform $M$ into a mixing matrix (in which the sums across each row equals one), we replaced the diagonal entries of $M_{kl}$ (within-county mixing) with one minus the sum of all entries outside county $k$:

$$M_{kk} \rightarrow 1 - \sum_{l \neq k} M_{kl}$$

Finally, we calculated the scaling factor $s$, such that the weighted proportion of sexual contacts within a county equals $\alpha$. We inferred $\alpha$ by fitting the model to the data:

$$\sum_{k} M_{kk} \frac{N_k}{\sum_k N_k} = \alpha$$

### Model parameterization

Model parameters are shown in Table 1 of the main text. We use data from Natsal-2[2] specifying the number of new sexual partners per year to parametrize parameters $c_j$ and $q_j$. We assumed that the distribution of reported number of new heterosexual partners is the sum of two Poisson distributions with means $c_j$, weighted by the proportion of individuals in each sexual activity class, $q_j$ (with $q_1 = 1 - q_2$).[3] Fitting to the data was performed with maximum likelihood estimation methods. The duration of asymptomatic untreated infection ($1/\gamma$) was taken from an evidence synthesis study.[4] We inferred the values of the variable parameters using Markov Chain Monte Carlo (MCMC) with a Metropolis Hasting algorithm, by comparing the time-dynamic model trajectories to the empirical data. First, we used data about the proportions of nvCT assuming a binomial likelihood. Second, we used CT diagnosis data assuming a negative binomial likelihood. We ran the model for 30 years to approach equilibrium in the absence of nvCT. We then simulated the emergence of nvCT in county C by assuming that 1% of prevalent CT cases in steady state change from wtCT to nvCT. We then set the time to Δ years before the discovery of nvCT and ran the model until 2015. We ran three separate MCMC chains each simulating 50.000 MCMC steps, using the R package BayesianTools. In the MCMC algorithm we assumed a binomial likelihood for the data about proportions of the nvCT. For the diagnoses data, we assumed a negative binomial likelihood with parameters $\mu(\theta)$ (the model computed number of diagnoses, as function of the model parameters $\theta$) and $var$ (the variance of number of diagnoses). The parametrization $NegBin(\mu, var)$ is sometimes referred to as the “ecological parametrization” of the negative binomial distribution [5], p. 165. It is a more dispersed distribution than the Poisson distribution through the factor $disp \geq 1$. If $disp = 1$ then this distribution is equal to the Poisson distribution. The relation with the original parametrization of the Negative Binomial distribution ($NegBin(r, p)$) is $r = \frac{\mu^2}{var-\mu}$ and $p = \frac{var-\mu}{var}$. So we take

$$var = \frac{\mu(\theta)^2}{\mu(\theta)*disp-\mu(\theta)} = \frac{\mu(\theta)}{disp-1}$$

The dispersion parameter $disp$ is not inferred in the MCMC sampling algorithm but, instead, a value of 10^0.5 is assumed for this parameter, reflecting a moderate dispersion. By this choice of dispersion, the data about proportions of the nvCT get a similar weight in the posterior log-likelihood as the diagnoses data.

Convergence of the MCMC chains was checked by computing the Gelman-Rubin convergence diagnostic.[6]

### S2: Data

In Tables S1 and S2 we have tabulated the data used in the MCMC simulations to infer the variable model parameters.

Table S1: Data used in the model on the proportion of the nvCT in different counties and years

| County | Number of samples with nvCT genotype/total number of samples | | | | |
|---|---|---|---|---|---|
| | 2006 | 2007 | 2008 | 2010 | 2014 |
| Blekinge | 7/106 | - | - | - | - |
| Dalarna | 520/812 | 104/204 | 42/172 | 64/253 | 18/292 |
| Halland | 140/584 | - | - | - | - |
| Kalmar | 38/188 | - | - | - | - |
| Norrbotten | 12/115 | 31/231 | 35/185 | 33/297 | 19/361 |
| Orebro | 63/162 | 97/261 | 55/233 | 34/151 | 25/374 |
| Södermanland | 36/119 | - | - | - | - |
| Skåne | 455/1896 | - | - | - | - |
| Stockholm | 26/115 | - | - | - | - |
| Uppsala | 50/263 | 62/230 | 37/206 | 31/262 | 18/336 |
| Västra Götaland | 24/93 | - | - | - | - |

Table S2: Data used in the model on the number of diagnoses in different counties and years

| County | 2004 | 2006 | 2007 | 2008 | 2009 |
|---|---|---|---|---|---|
| Dalarna | 1037 | 907 | 2446 | 1579 | 1293 |
| Norrbotten | 964 | 1023 | 967 | 991 | 969 |
| Orebro | 875 | 854 | 1294 | 1245 | 1282 |
| Uppsala | 1194 | 1353 | 1611 | 1490 | 1218 |

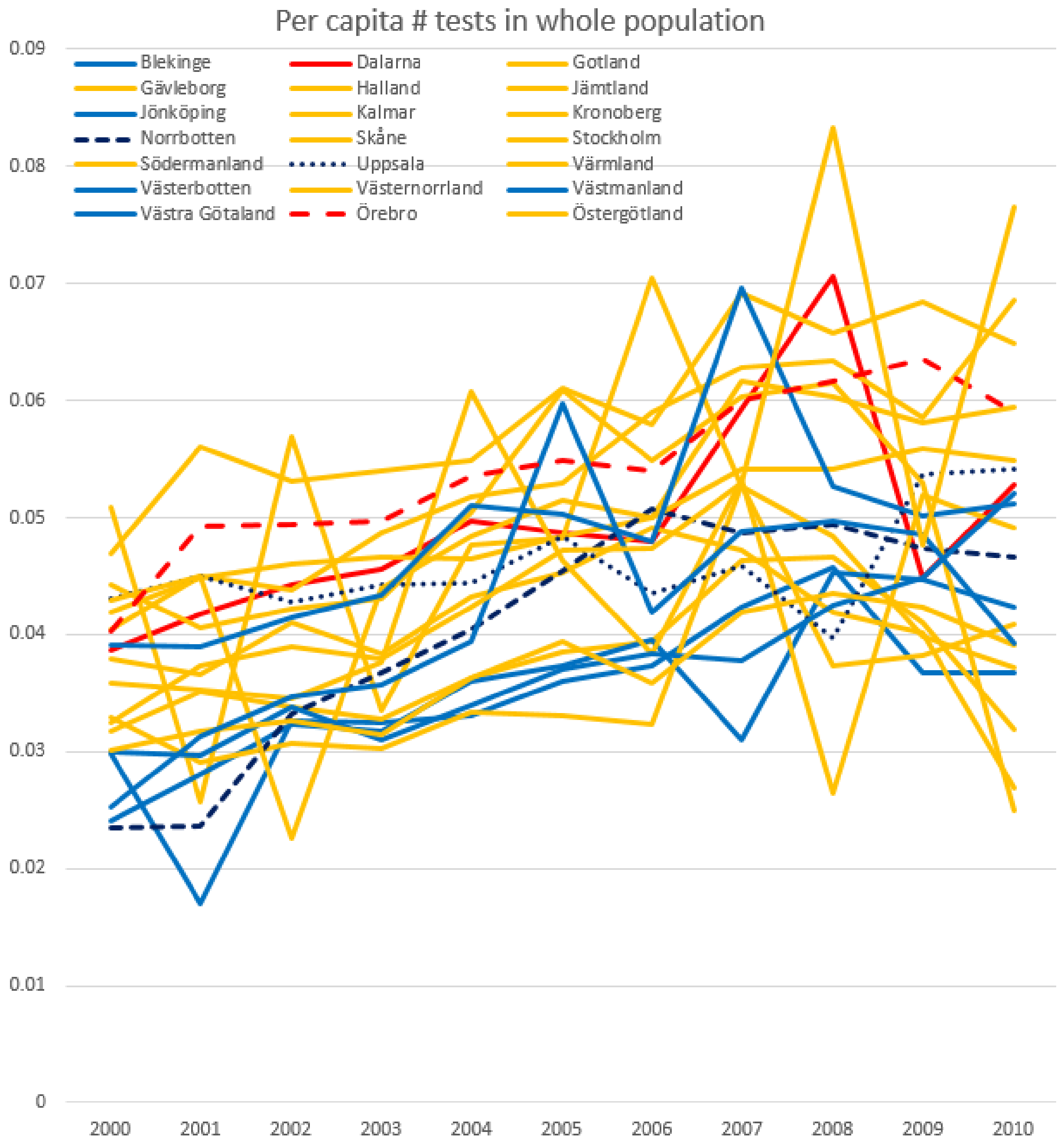

Figure S2: Per capita number of tests in the whole population (data not used in the model)

### S3: Basic reproduction rates

The basic reproduction number, $R_0$, can be calculated using the next-generation matrix method as described by Diekmann et al.[7,8] It is different for the wtCT, the nvCT before discovery and the nvCT after discovery. As we did not consider sex-specific differences in sexual behavior or the natural history of chlamydia, and assumed that the sexual behavior of individuals is the same across all Swedish counties, we can simplify the model into a single population with two different sexual activity groups. For wtCT, the transmission matrix $T$ is given by

$$T = \begin{bmatrix} \beta c_l \rho_{ll} N_l / N_l & \beta c_l \rho_{lh} N_l / N_h \\ \beta c_h \rho_{hl} N_h / N_l & \beta c_h \rho_{hl} N_h / N_h \end{bmatrix}$$

and contains the infection rates of individuals in the two sexual activity classes, stratified by the activity class of the sexual partner. $\rho$ corresponds to the mixing matrix between sexual activity classes as described above.

For nvCT, the transmission matrix is equal before and after discovery. It is given by $fT$, where $f$ is the difference in biological fitness between wtCT and nvCT.

The transition matrix $\Sigma$ for wtCT and nvCT after discovery is given by

$$\Sigma = \begin{bmatrix} -(\gamma + \tau) & 0 \\ 0 & -(\gamma + \tau) \end{bmatrix}$$

The transition matrix $\Sigma$ for nvCT before discovery is given by

$$\Sigma = \begin{bmatrix} -\gamma & 0 \\ 0 & -\gamma \end{bmatrix}$$

$R_0$ is defined as the dominant eigenvalue of the next-generation matrix $K = -T\,\Sigma^{-1}$.

### S4: Sensitivity analysis: biological fitness affects duration of infection

Table S1 shows the prior and posterior distributions of the model parameters if we assumed that a difference in biological fitness between the wtCT and the nvCT affects the duration of infection, rather than the per partnership transmission probability (Table 1 main text). For the nvCT, parameter $\gamma$ is replaced by $\gamma / f$ in the model equations.

Table S3: Description of fixed and variable model parameters, and model derived quantities

| | Fixed parameters | Value | Source |
|---|---|---|---|
| $q_j$ | Proportion in risk group | 0.93, 0.07 | Ref 2 |
| $c_j$ | Partner change rates | 0.59, 6.57 | Ref 2 |
| $\gamma$ | Chlamydia clearance rate (per year) | 0.73 | Ref 4 |
| $C$ | County where nvCT emerged | 1-21 | Methods |
| | **Variable parameters** | **Prior** | **Posterior*** |
| $\rho$ | Between-county mixing dependency on distance | unif(0,2) | 1.12 (0.87-1.34) |
| $\alpha$ | Average fraction of new contacts inside of county | unif(0,1) | 0.75 (0.72-0.77) |
| $\beta$ | Per partnership transmission probability | unif(0,1) | 0.93 (0.84-1.00) |
| $\epsilon$ | Assortativity index risk groups | unif(0,1) | 0.04 (0.00-0.16) |
| $f$ | Relative fitness of nvCT compared to wtCT (affects duration of infection) | unif(0.7,1.3) | 0.18 (0.16-0.19) |
| $\tau$ | Treatment rate (wtCT) per year | unif(0,3) | 2.31 (1.97-2.70) |
| $\pi$ | Maximal increase of treatment rate $\tau$ after October 2006 | unif(0,0.25) | 0.00 (0.00-0.02) |
| $\Delta$ | Number of months that the nvCT remained undiscovered until October 2006 | unif(36,144) | 44 (37-52) |
| | **Model derived quantities** | | **Posterior*** |
| | Year of emergence | | Jan '03 (Nov '02-Aug '03) |
| | Proportion treated** | | 0.76 (0.73-0.79) |
| | Prevalence before emergence | | 1.03 (0.85-1.21) |
| | Max prevalence*** | | 3.01 (2.62-3.55) |
| | $R_0$ wtCT | | 1.07 (1.06-1.08) |
| | $R_0$ nvCT before discovery | | 3.66 (3.27-4.08) |
| | $R_0$ nvCT after discovery | | 1.01 (1.00-1.02) |

* median (95% credible interval). Only results for the model in which we assumed that nvCT emerged in Dalarna (M1) are shown. Assumed: difference in biological fitness between wtCT and nvCT affects duration of infection.

** Computed as $\tau/(\tau+\gamma)$.

*** Prevalence in Dalarna, October 2006.

### S5: Proportion of infected people that is treated

We found a posterior distribution for the (wtCT) treatment rate $\tau$ that implies that 75% (72-77%) of infected people is treated (Table 1 main text). We cannot use the model itself to verify whether this is a realistic finding, because the model does not include explicitly all processes by which people are treated, including asymptomatic screening. To further analyze how credible the posterior distribution of the treatment rate is, we do the following (back-of-the-envelope) calculations:

Infected people can become susceptible again by natural recovery (at rate $\gamma$), by receiving treatment for symptoms (at rate $\xi$, not considered in the model) or by asymptomatic screening (at rate $\sigma$, not considered in the model). Probably, asymptomatic screening rates are higher in infected compared to susceptible people, because infected people tend to have more risky sexual behavior. Screening campaigns may therefore be more targeted towards infected people, and infected people are more likely to be screening through notified, infected partners. We define $\eta$ as the ratio of screening rates in infected compared to that in susceptible people. Then, the fraction of the population that is screened per year is:

$$f_{screened} = (1 - prev) * \left(1 - e^{-\sigma/\eta}\right) + prev * (1 - e^{-\sigma})$$

We solve this equation for $\sigma$. We have data on the fraction of the population that is screened per year (0.25)[9] and the (model-computed) prevalence in the population aged 15-29 that is considered in the model (0.01). However, $\eta$ is not well known, so we make assumptions about $\eta$ instead (we consider a range of values between 1 and 10).

Further, suppose that there was no screening. Then only symptomatically infected people would be treated:

$$f_{symp} = \frac{\xi}{\xi + \gamma}$$

We can also solve for $\xi$, using $\gamma$=0.73 [4] and considering a range of values between 0 and 1 for $f_{symp}$. Then we compute the proportion of infections that is treated as:

$$f_{treat} = \frac{\sigma + \xi}{\sigma + \xi + \gamma}$$

We then verify which values for $\eta$ and $f_{symp}$ values are consistent with $f_{treat}$ between 0.72 and 0.77 (red area in Figure S3). These computations show that if $\eta$ is around 5, and $f_{symp}$

between 30-60%, the model-computed value for $f_{treat}$ is not unlikely. We deem such values for $\eta$ and $f_{symp}$ in a credible range.[10,11]

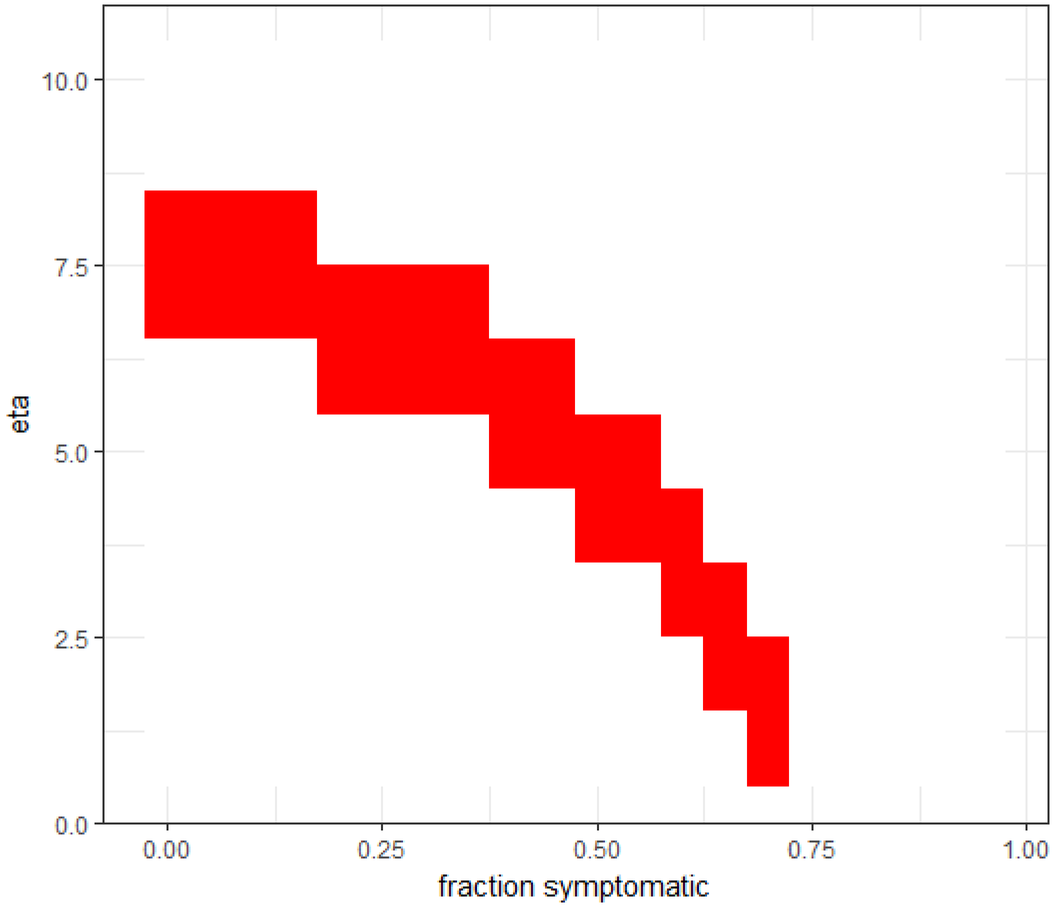

Figure S3: Red area: values for $\eta$ (ratio of screening rates in infected compared to susceptible people) and $f_{symp}$ (proportion symptomatic) when the prevalence is 0.01 and the proportion of infected people that is treated is between 0.72 and 0.77.
